# Sensitive and specific spectral library searching with COSS and Percolator

**DOI:** 10.1101/2021.04.09.438700

**Authors:** Genet Abay Shiferaw, Ralf Gabriels, Robbin Bouwmeester, Tim Van Den Bossche, Elien Vandermarliere, Lennart Martens, Pieter-Jan Volders

## Abstract

Maintaining high sensitivity while limiting false positives is a key challenge in peptide identification from mass spectrometry data. Here, we therefore investigate the effects of integrating the machine learning-based post-processor Percolator into our spectral library searching tool COSS. To evaluate the effects of this post-processing, we have used forty data sets from two different projects and have searched these against the NIST and MassIVE spectral libraries. The searching is carried out using two spectral library search tools, COSS and MSPepSearch with and without Percolator post-processing, and using sequence database search engine MS-GF+ as a baseline comparator. The addition of the Percolator rescoring step to COSS is effective and results in a substantial improvement in sensitivity and specificity of the identifications. COSS is freely available as open source under the permissive Apache2 license, and binaries and source code are found at https://github.com/compomics/COSS

## Introduction

MS-based peptide identification typically relies on matching measured spectra against theoretical spectra in a database searching approach^1^. However, identification can also be obtained by matching measured spectra against a spectral library consisting of previously measured and identified spectra^2^. Several spectral library searching tools have been developed for this purpose, with notable examples including SpectraST^3^, the National Institute of Standards and Technology (NIST) MS Search^4^ and MSPepSearch (https://chemdata.nist.gov/dokuwiki/doku.php?id=peptidew:mspepsearch), ANN-SoLo^5^, X!Hunter^6^, and COSS^7^. The direct comparison of a newly measured spectrum against the spectra in such a spectral library can both increase sensitivity^8^ and reduce computational complexity compared to database searching^9^.

However, similar to database searching, spectral library searching is not perfect, and several causes can lead to incorrect peptide identification^10^: quality of the spectral data, unexpected post translational modifications, charge state issues, or a poor scoring function. To control this erroneous identification, the validation of search results is a crucial step in the identification process. Typically, this is handled through a target-decoy approach in both database and spectral library searches^1^. This approach allows estimation of the False Discovery Rate (FDR)^11,12^, which is the expected proportion of incorrect peptide to spectrum matches (PSMs) among the selected set of accepted identifications.

Besides the validation of PSMs using target-decoy based FDR control, it has also become common to employ post-processing methods to database search results to increase sensitivity. The most popular of these is Percolator^13^, which significantly improves the sensitivity of multiple database search engines, including SEQUEST, Mascot^14^, and MS-GF+^15^. Percolator itself is a semi-supervised machine learning algorithm based on a linear support vector machine, which is designed to discriminate between correct and incorrect peptide matches by rescoring peptide identifications. For this, Percolator considers a set of features that describe each PSM and uses these features as well as the annotation of target and decoy PSMs in an iterative process to re-rank PSMs using a new score and associated q-value. Yet, although Percolator is commonly used in database^16,17^ searches, its use has thus far not been reported for spectral library searching.

Here, we have therefore examined the utility of Percolator in spectral library searching by extending our COSS spectral library search tool. Importantly, we have found that Percolator improves the output of COSS in two ways. First, Percolator rescoring increases the total number of identifications obtained, thus improving sensitivity. Second, Percolator use also adds more reliable PSMs and deletes false positive PSMs from the original search result, thus improving specificity as well.

We therefore integrated Percolator into the latest version of our freely available, open-source COSS tool, which is moreover capable of handling multiple file formats, and can also be used to analyze large data sets.

## MATERIAL AND METHODS

### Experimental data sets and spectral library

We obtained raw data files from the deep proteome and transcriptome abundance atlas^18^ data set (36 runs corresponding to one fractionated brain sample; ProteomeXchange ID PXD010154) and four runs from “Assessing the relationship between mass window width and retention time scheduling on protein coverage for data-independent acquisition”^19^ data set (ProteomeXchange ID PXD013477, DDA runs of the HeLa sample) as benchmarking data sets (Supplementary Table S-1). All these raw files were converted to Mascot Generic Format (mgf) format using the msconvert tool (ProteoWizard^20^ version 3.0.19014), with the peak picking algorithm activated. The well-known NIST spectral library (https://chemdata.nist.gov/dokuwiki/doku.php?id=peptidew:cdownload, obtained on 15/08/2021_) and MassIVE^21^ (obtained on 18/09/2018) were used to perform our searches against.

### Spectral library searches

All 40 data sets are searched with COSS against the NIST and MassIVE spectral library, which were appended with the corresponding decoy spectra. The following search settings were used: precursor mass tolerance set to 10 ppm, fragment mass tolerance set to 0.05 Dalton (Da), MSROBIN scoring function^7^, and a fragment mass window to select peaks set to 10 Da.

To run MSPepSearch (version 0.96), we first generated and concatenated an MSPepSearch decoy spectral library using the COSS decoy generator. Next, we converted the msp file of this library to MSPepSearch’s binary file format using Lib2NIST (version 1.0.6.5) and performed the searches with a precursor mass tolerance of 10 ppm and fragment mass tolerance 0.05 Da.

### Sequence database search

In parallel with the spectral library searches, we performed a sequence database search with the well-established MS-GF+^22^ (version 2021.01.08) tool through SearchGUI^23^ (version 4.0.22) with additional features reporting activated for use by Percolator. The search database was constructed from the human reference proteome (UP000005640) as obtained from UniProtKB^24^ (consulted on 9/10/2018). Carbamidomethylation of cysteine was set as a fixed, and oxidation of methionine as a variable modification. Trypsin was set as protease and a maximum of two missed cleavages was allowed. Precursor mass tolerance was set to 10 ppm, and fragment mass tolerance to 0.05 Da. Precursor charges from 2 to 4 were considered. The SearchGUI configuration file used to run MS-GF+ is provided in Supplementary methods.

### Percolator rescoring

We integrated Percolator^13^ version v3-04 into COSS^7^ such that Percolator’s input features are generated from COSS results and Percolator can be executed automatically. COSS implements multiple scoring functions, and all scores as well as additional matching peak features are exported as features to Percolator to maximize the amount of information available for rescoring. All features with descriptions are found in Supplementary Table S-2. These features are output to a tab-delimited text file which Percolator can use automatically.

For MSPepSearch, Percolator was provided with the full list of available features from the output provided, and these are also found in Supplementary Table S-2. In the case of MS-GF+, the Percolator input text file is generated using the msgf2pin command provided by MS-GF+^15^.

Percolator was run with the target-decoy competition method enforced as all searches are performed against a concatenated database. In addition, the error check is overridden to ensure the output contains the rescored q-values as obtained from the Support Vector Machine (SVM). Identifications from the fractionated sample were concatenated prior to rescoring.

### Retention time prediction

In order to evaluate how rescoring with Percolator improves true positive identifications, we used an orthogonal validation which makes use of DeepLC^25^ (version 0.1.35), a deep learning algorithm that accurately predicts peptide retention times. Input files for DeepLC were generated from the search results using custom scripts. In addition, DeepLC calibration files were constructed by selecting the 1000 peptides with the highest score across ten equally sized retention time windows.

## RESULTS AND DISCUSSION

### Effects of different feature sets on the rescored result

Percolator uses a machine learning algorithm to rescore the output of peptide identification tools to achieve higher sensitivity and specificity. The result of this process depends strongly on the quality and utility of the input features. Here we therefore supplemented standard COSS output with additional scores and features for each PSM to maximize the potential of the rescoring step. Principal component analysis (Supplementary Figure S-1) shows several distinct clusters in the features, indicating that each group of features has the potential to add unique information to the SVM trained by Percolator. To examine the effect of these features on rescoring in spectral library searching, we analyzed different combinations of input features with Percolator. The full set of inputs to Percolator and their description can be found in Supplementary Table S-2. Figure 1 shows that using all features from COSS yields the best results in terms of identification rate at 1% FDR (28.9%). While removal of precursor mass and charge has little effect on the identification rate, the intermediate scores generated by COSS (peak and intensity fraction of matched spectra) does lower the identification rate substantially (15.9%).

**Figure 1.**
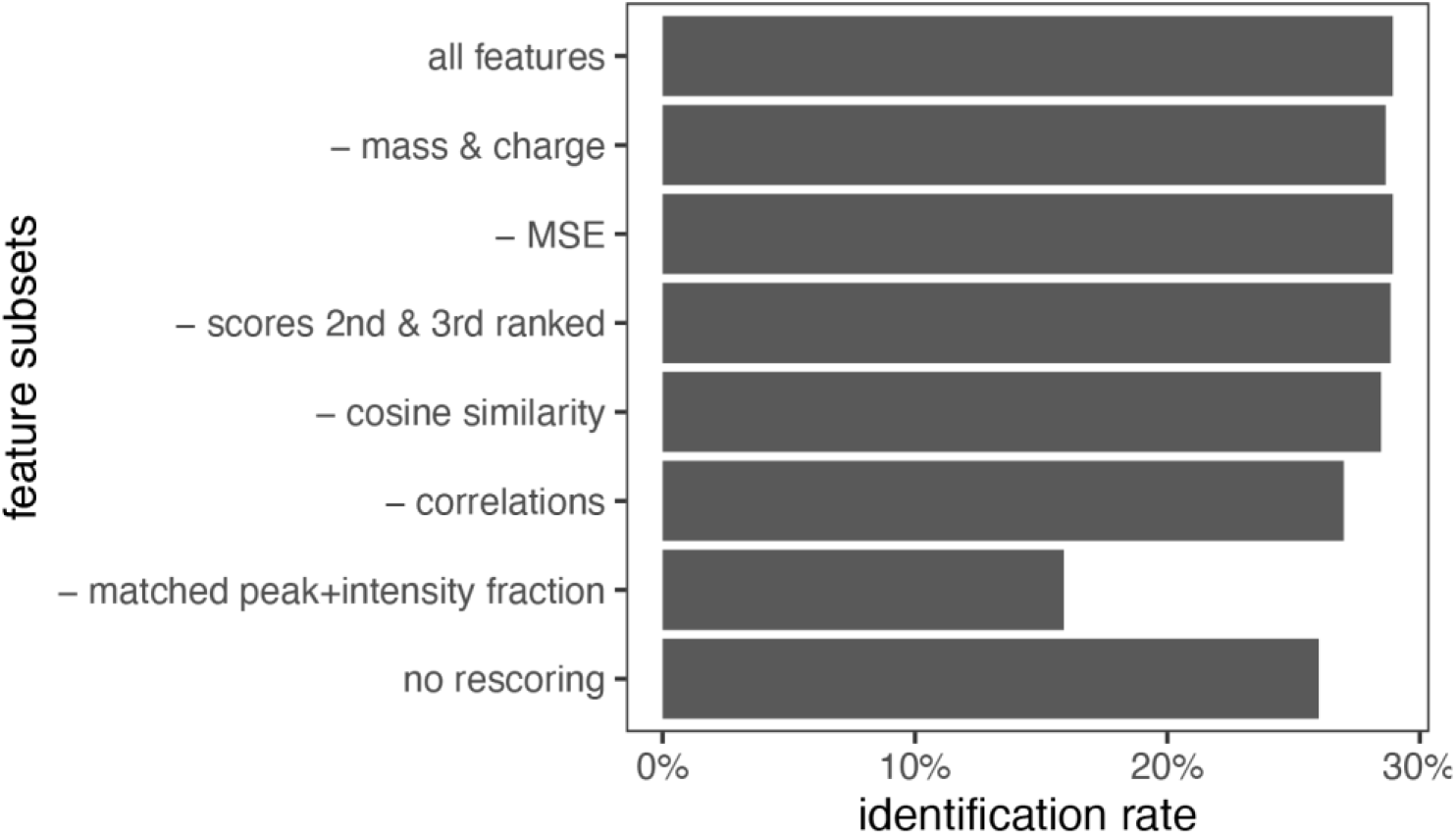
Effect of different feature sets provided to Percolator as measured by identification rate at 1% FDR.

### Comparison of COSS, MSPepSearch and MS-GF+ with and without Percolator

To put the improvement observed in rescoring COSS results into context, we compared these with rescoring results from MSPepSearch, a different but performant spectral library search tool, and MS-GF+, a popular database search engine. Figure 2 shows the identification rates obtained either at 1% FDR (without Percolator), or for a q-value at or below 0.01 (with Percolator) for each of the two test data sets as analyzed by COSS, MSPepSearch, and MS-GF+. For all datasets, the COSS search against MassIVE consistently outperforms both MSPepSearch and MS-GF+, and this for both pre- and post-Percolator identification results. In the case of data sets from PXD010154, MS-GF+ has slightly more identifications than COSS against the NIST spectral library, which can likely be attributed to incomplete coverage of the library for this sample. This effect is also reflected in the even lower performance of MSPepSearch on this data set. In the case of PXD013477, identification is quite high for all three tools, in line with the results in the original manuscript^19^. Overall, these results show that rescoring the output of spectral library search engines can drastically increase identification rate. Depending on the coverage of the spectral library, the identification rate can exceed those obtained with database search engines.

**Figure 2.**
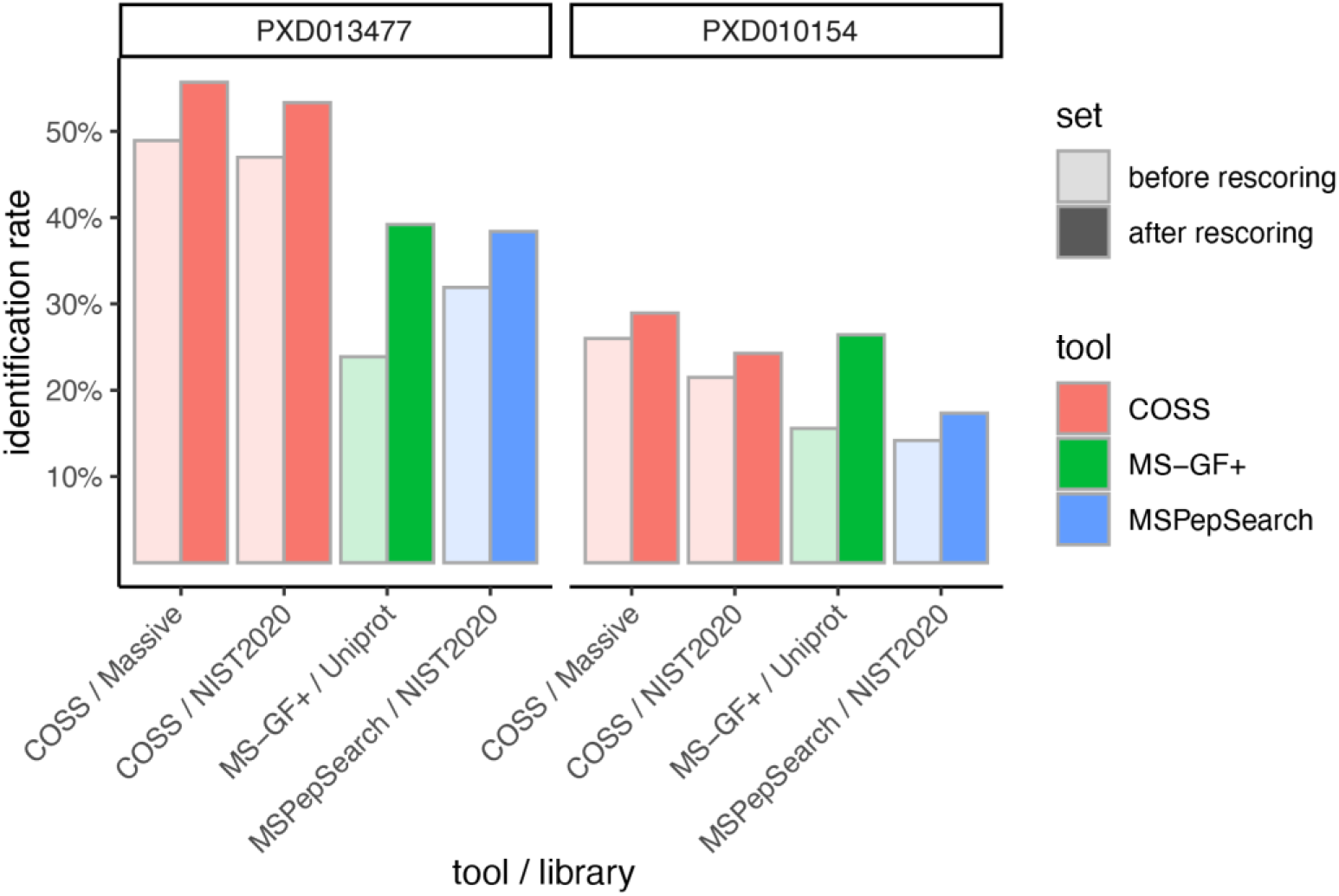
Comparison of identification rates achieved by COSS, MSPepSearch and MS-GF+, before rescoring and after rescoring. All results are either taken at 1% FDR (before rescoring) or at a q-value cut-off at 0.01 (after rescoring).

### Comparison of obtained identifications before and after Percolator rescoring

To gain further insight into the difference between results obtained with, and without Percolator rescoring, the overlap at 1% FDR before and after rescoring was analyzed at the peptide level. Figure 3 shows this identification agreement for COSS with and without Percolator. For all data sets, Percolator removes some of the identified peptides (PXD013477: 3%; PXD010154: 3.4%), while adding a larger set of new results (PXD013477: 16%; PXD010154: 13%). The vast majority (PXD013477: 97%; PXD010154: 96.6%) of the identifications is maintained, however. Upon comparing the length of the peptides added and removed by Percolator, it appears that Percolator improves the sensitivity for small peptides (median length of ten amino acids), while removing preferentially larger peptides (Figure S-2). No significant difference in the amino acid composition of added *versus* removed peptides can be observed (Figure S-3).

**Figure 3.**
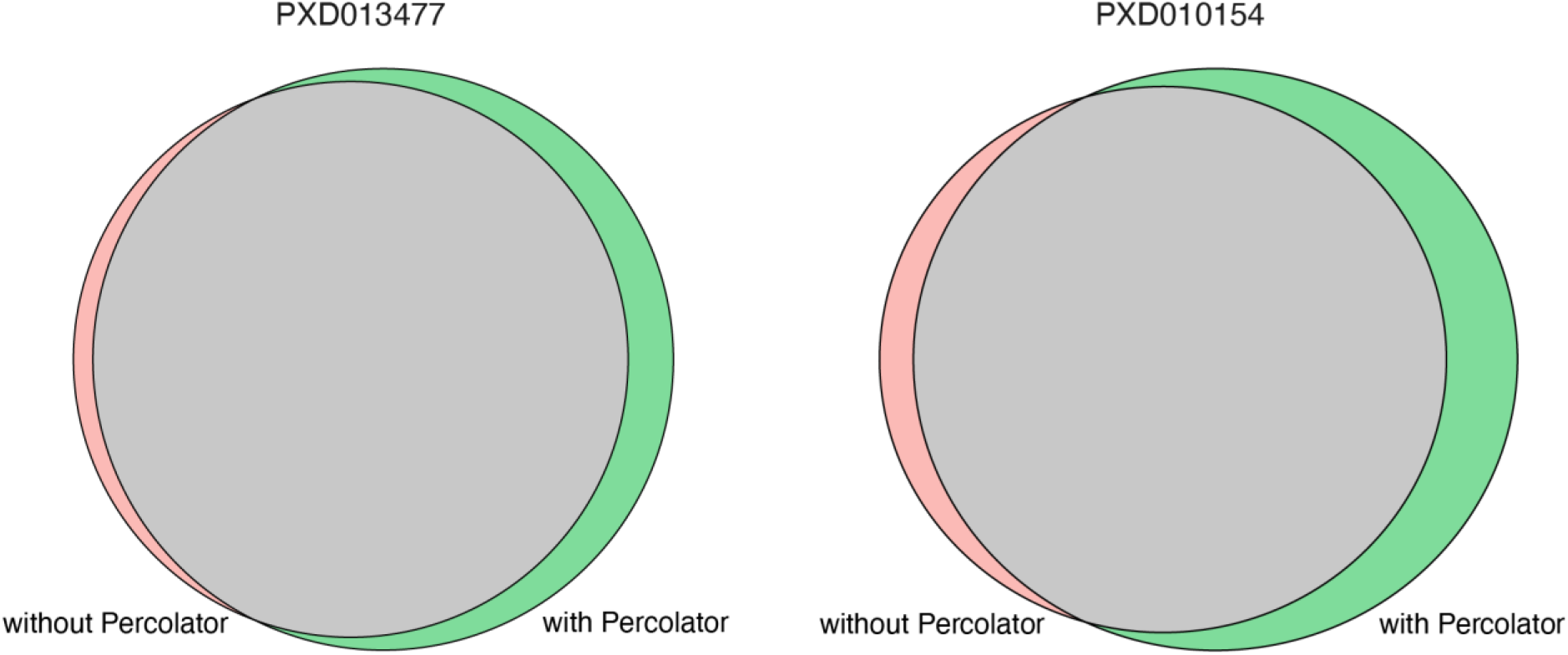
Peptide level overlap in search results from COSS against the NIST library at the 1% FDR level before and after Percolator rescoring. The number of peptides added by rescoring exceeds the number of removed peptides, resulting in an overall increase of the identification rate.

### Orthogonal validation of the rescoring results using measured and predicted retention time

With the drastically increased reliability of retention time prediction, the comparison of the observed retention time and the predicted retention time for a specific peptide has been put forward as a means of orthogonal validation for PSMs^26,27^. Here, an overall poor correlation of predicted and observed retention time is indicative of a misidentified peptide. We therefore performed retention time prediction using DeepLC and compared the predicted with the observed retention time for PSMs affected by Percolator (Figure 4). The identifications added by rescoring had an overall smaller absolute difference in predicted *versus* observed retention time than the identifications removed by rescoring. This indicates that the latter set contains more misidentifications that are correctly removed by Percolator. In addition, the distribution of the difference in predicted *versus* observed retention time of Percolator added identifications closely resembles the distribution of the unchanged identifications, suggesting that these are indeed valid identifications.

**Figure 4.**
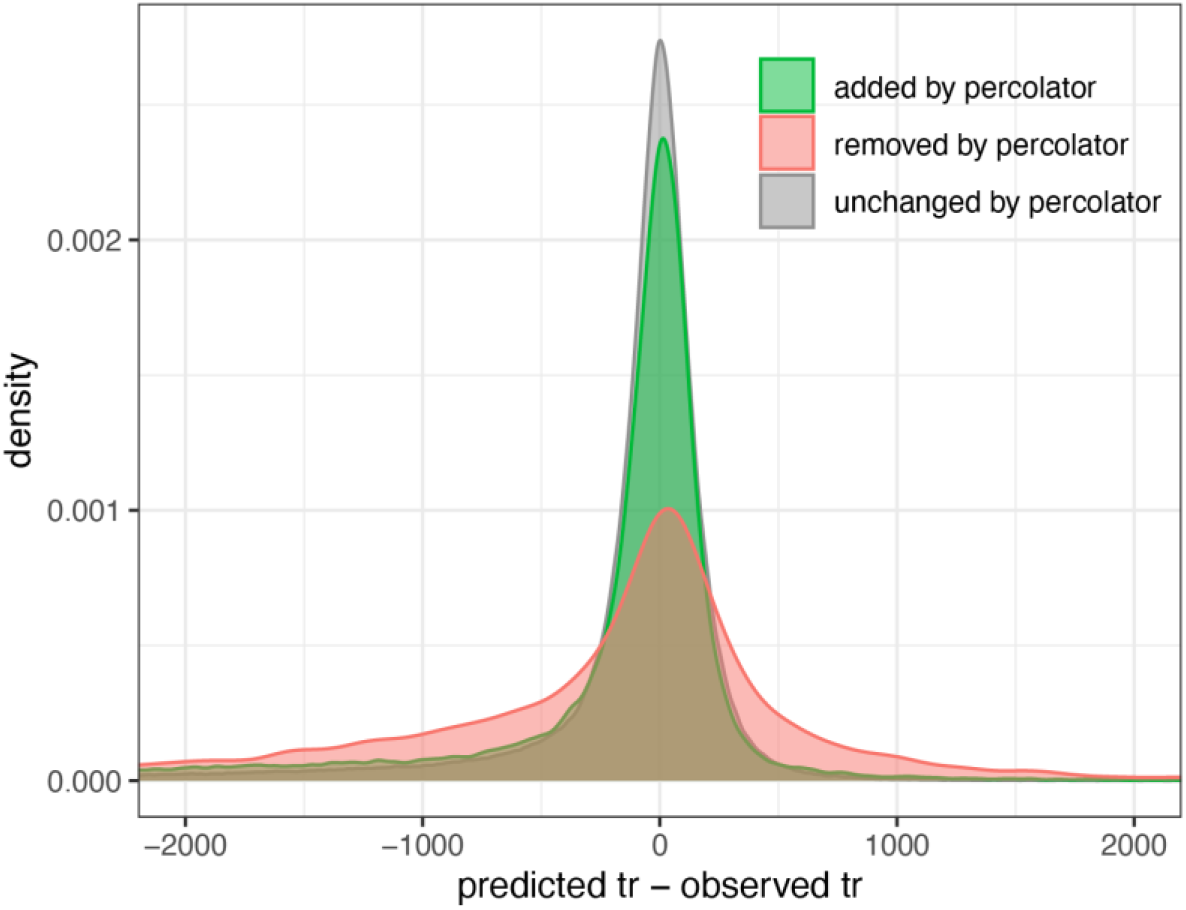
Comparison of measured and predicted retention time of peptide identification added and removed by rescoring. Shown here are identifications obtain with COSS on the PXD010154 data set and the NIST spectral library. The high resemblance between the distribution for the added and unchanged identifications suggests that there are indeed valid PSMs while the discrepancy with the distribution of the remove identifications indicates that those are enriched with misidentifications.

## CONCLUSION

Increasing sensitivity while maintaining specificity is key in peptide and protein identification from mass spectrometry data. Rescoring peptide identifications using tools such as Percolator to increase sensitivity and specificity is thus common practice in sequence database searching. However, until now it was unclear if such post-processing had any benefits for spectral library searching. Here, we have shown that combining Percolator with such tools enhances sensitivity, and that it can also enhance specificity. Specifically, our COSS spectral library search tool shows increased sensitivity, and dramatically enhanced specificity when combined with Percolator. We have thus shown that, for COSS at least, the combination of these benefits justifies the added complexity of the percolator post-processing step. Based on these findings, we can thus recommend use of rescoring in spectral library searching. To this end, the latest version of COSS (COSS-2.0) has been fitted with Percolator integration.

## AVAILABILITY

The COSS software and its source code can be freely downloaded from https://github.com/compomics/COSS and is licensed under the permissive, open-source Apache License, version 2.0.

## ACKNOWLEDGMENTS

This project is supported by the National Institute of Health (NIH) [NCI-ITCR grant number 1U24CA199347 to G.A.S.], Research Foundation - Flanders (FWO) [grant number 1S50918N to R.G., grant number 1S90918N to T.V.D.B., G042518N and G028821N to L.M., and grant number 1253321N to P.V.], by the European Union’s Horizon 2020 Program (H2020-INFRAIA-2018-1) [823839 to L.M.], and Ghent University Concerted Research Action [grant number BOF21-GOA-033 to L.M.]. R.B. acknowledges funding from the Vlaams Agentschap Innoveren en Ondernemen under project number HBC.2020.2205. The authors would like to thank all CompOmics group members for their ideas, discussions, and support.

